# A circulating biomarker of facioscapulohumeral muscular dystrophy clinical severity, valid in skeletal muscle and blood

**DOI:** 10.1101/2022.08.31.506017

**Authors:** Christopher R. S. Banerji, Anna Greco, Leo A. B. Joosten, Baziel van Engelen, Peter S. Zammit

**Affiliations:** King’s College London, Randall Centre for Cell and Molecular Biophysics, New Hunt’s House, Guy’s Campus, London SE1 1UL, UK; Department of Neurology, Donders Institute for Brain, Cognition and Behaviour, Radboud University Medical Center, Nijmegen, The Netherlands; Department of Internal Medicine, Radboud Institute of Molecular Life Sciences (RIMLS) and Radboud Center of Infectious Diseases (RCI), Radboud University Medical Center, Geert Grooteplein Zuid 10, 6525 GA Nijmegen, the Netherlands; Department of Medical Genetics, Iuliu Hatieganu University if Medicine and Pharmacy, Cluj-Napoca, Romania

**Keywords:** Facioscapulohumeral muscular dystrophy, FSHD, circulating biomarker, transcriptomics, MRI, DUX4, PAX7

## Abstract

Facioscapulohumeral muscular dystrophy (FSHD) is incurable. DUX4 mis-expression is believed to underlie FSHD pathogenesis, alongside PAX7 target gene repression, yet clinical trials lack robust biomarkers of severity. FSHD entails fatty replacement of muscle, accelerated by inflammation, we thus performed RNA-sequencing on both an MRI guided inflamed (TIRM+) and non-inflamed (TIRM-) muscle biopsies from clinically-characterised FSHD patients, alongside peripheral blood mononucleated cells (PBMCs). PAX7 target gene repression in TIRM-muscle associates with severity. DUX4 target gene biomarkers associate with lower limb fat fraction and D4Z4 repeat length, but not severity. PAX7 target gene repression in muscle correlates with levels in matched PBMCs. A refined biomarker computed in PBMCs associates with severity in FSHD patients, and also validates as a classifier of severity in an independent set of 54 FSHD patient blood samples. In summary, we present a minimally-invasive, circulating, transcriptomic biomarker of FSHD clinical severity valid in muscle and blood.

## Introduction

Facioscapulohumeral muscular dystrophy (FSHD) is a prevalent (∼12/100,000^1^), incurable, inherited myopathy. FSHD is slowly progressive and highly heterogeneous both clinically and molecularly^2^, making reliable biomarkers for clinical severity and progression challenging to develop. However, such biomarkers are needed for monitoring FSHD patients and measuring outcomes in clinical trials. Moreover, gene-based biomarkers facilitate understanding of pathomechanisms and inform therapeutic development.

FSHD is linked to epigenetic derepression of the D4Z4 macrosatellite at chromosome 4q35 alongside a permissive 4qA haplotype. Epigenetic derepression is achieved by two mechanisms. The most common (95% of patients, FSHD1, OMIM: 158900) requires truncation of the D4Z4 region from the typical >100 repeats to 10-1 units^3^. Mutations in epigenetic modifiers, typically *SMCHD1*^4^ but rarely *DNMT3B*^5^ or *LRIF1*^6^ drive epigenetic derepression in the remaining 5% (FSHD2, OMIM: 158901). Each D4Z4 repeat encodes a transcription factor termed DUX4 and epigenetic derepression permits transcription of DUX4 from the distal-most D4Z4 unit. DUX4 transcripts are stabilised by splicing to a poly(A) signal in the flanking DNA on permissive 4qA haplotypes, permitting translation^3^.

DUX4 is expressed during zygotic genome activation, before being epigenetically repressed in somatic tissues^7^. Mis-expression of DUX4 is believed to underlie FSHD pathogenesis, but is extremely difficult to detect in FSHD muscle, with protein found in ∼1/1000 FSHD myoblasts ex vivo^2,8^. Despite this, DUX4 target gene biomarkers associate with FSHD status, particularly in the context of active inflammation (e.g., STIR/TIRM positivity on MRI)^9–14^. The two DNA-binding homeodomains of DUX4 show homology to the single homeodomain of the myogenic master regulator PAX7, suggesting inhibitory interactions between DUX4 and PAX7^10,15^. We have demonstrated that PAX7 target gene repression hallmarks FSHD muscle regardless of inflammatory state, associating with histological and MRI measures of pathological severity^9–11^. Our PAX7 target gene repression biomarker has been independently verified^14,16^. Importantly, PAX7 target gene repression progresses in muscle biopsies from the same FSHD patient one year apart, associating with disease duration^17^.

Therapeutics for suppressing DUX4 have reached clinical trials, including the p38 inhibitor losmapimod^18^. However, losmapimod induced no significant change in the primary outcome measure: DUX4 target gene expression in FSHD muscle, despite improvement in secondary functional outcomes. This motivates development of more sensitive biomarkers of FSHD severity. However, biomarkers based on muscle biopsies are invasive, and FSHD muscle may not mount an appropriate regenerative response, necessitating minimally invasive measures^19^.

Circulating biomarkers for FSHD have been investigated via discovery approaches. Proteomics of muscle microdialysates identified innate immunity mediators S100-A8 and A9^20^ as FSHD associated. Studies of FSHD plasma also found S100-A8, several miRNAs^21^ and complement components^22^. We defined a set of genes over-expressed in FSHD lymphoblastoid cell lines, which correlate with immune infiltration in FSHD muscle biopsies^11^. Recently, a large RNA-sequencing study on peripheral blood found no transcripts differentially expressed between FSHD patients and controls, or associated with FSHD clinical severity^23^. Conversely, serological studies identified IL-6^24^, TNFα^25^ and structural muscle components^26^ as associated with FSHD severity: how the circulating levels of these factors impact FSHD muscle tissue is unknown.

FSHD is a muscle pathology, accelerated by inflammation^2^ and informative blood borne biomarkers should correlate directly to muscle-based markers of severity. To identify biomarkers of clinical severity valid in both tissues, we performed RNA-sequencing on 24 clinically characterised FSHD patients, obtaining paired inflamed (TIRM+) and non-inflamed (TIRM-) muscle biopsies and peripheral blood mononucleated cells (PBMCs), alongside muscle and PBMCs from control subjects. We computed DUX4 and PAX7 biomarkers and identified associations between biomarker levels and measures of clinical severity. There was clear association between levels of PAX7 target gene repression in muscle and blood, but not DUX4 target gene expression. Importantly, a refined 143 PAX7 target gene signature (the ‘FSHD muscle-blood biomarker’) in PBMCs correlated with widely used FSHD clinical severity scores defined by Ricci^27^ and Lamperti^28^. Investigation of our FSHD muscle-blood biomarker on an independent dataset of 54 FSHD and 29 matched control blood samples validated it as a minimally invasive biomarker of FSHD severity, with particular relevance in older patients.

## Results

### Clinical assessments correlate strongly in FSHD

49 individuals were investigated: 25 FSHD1 patients, 1 FSHD2 patient and 23 control individuals. Clinical variables obtained included: age, sex and MRC sum score for 12 muscle groups. For FSHD patients severity indicators were also measured including: D4Z4 repeat length (for FSHD1 patients), Ricci clinical severity score^27^, Lamperti clinical severity score^29^, disease duration, lower limb fat fraction (LLFF) assessed by MRI and maximum voluntary contraction (MVC) of tibialis anterior (TA) (**Table S1**). Considering the entire cohort, there were no significant differences in sex (logit regression *p=*0.31, FSHD 46% male, control 61% male, **Table 1**), however age at examination was significantly higher in FSHD patients versus controls (Wilcoxon *p*=2.9×10^−3^, **Table 1**). Total MRC sum score was higher in controls (all scored 60/60, Wilcoxon *p*=3.0×10^−9^, **Table 1)**.

**Table 1:** Summary of clinical variables associated with control and FSHD patients. Table presents sample counts, age at examination, % males, MRC Muscle grade for 12 muscle groups, D4Z4 repeat length (for FSHD1 patients), Ricci clinical severity score, Lamperti clinical severity score, Disease duration, LLFF and MVC of TA, separately for all samples and those samples undergoing muscle biopsy and PBMC isolation. Mean values for each variable are presented alongside the range in brackets. For age at examination and MRC Muscle grade a *p*-value represents the significance of a Wilcoxon test comparing control and FSHD affected individuals in the 3 groups. For % male a *p*-value represents the significance of logit regression comparing the sex distribution in control and FSHD affected individuals across the 3 groups.

|  | All Samples |  |  | Muscle Biopsies |  |  |
| --- | --- | --- | --- | --- | --- | --- |
|  | Control | FSHD | *p*- value | Control | FSHD | *p*- value |
| n | 23 | 26 | - | 11 | 24 | - |
| Age at examination (years) | 38.3 (21,65) | 50.3 (30,63) | 2.90E-03 | 31.6 (21,63) | 49.8(30,63) | 5.20E-03 |
| Sex %M | 60.8 | 46.1 | 0.31 | 81.8 | 45.8 | 0.047 |
| MRC Muscle grade (max 60) | 60 (60,60) | 51.6 (26,60) | 3.04E-09 | 60 (60,60) | 51.25 (26,60) | 1.10E-05 |
| D4Z4 repeat length | - | 6.96 (3,9) | - | - | 7.0 (3,9) | - |
| Ricci Score (max 10) | - | 4.92 (2,8) | - | - | 4.96 (2,8) | - |
| Lamperti Score (max 15) | - | 5.81 (1,11) | - | - | 6 (1,11) | - |
| Disease Duration | - | 24 (2,52) | - | - | 24.4 (2,52) | - |
| Lower Limb fat fraction (%) | - | 14.0 (2.9,56.3) | - | - | 14.4 (2.9,56.3) | - |
| MVC of tibialis anterior (N) | - | 69.6 (9.2,201.0) | - | - | 69.8 (9.2,201.0) | - |
|  | PBMCs |  |  |  |  |  |
|  | Control | FSHD | *p*- value |  |  |  |
| n | 14 | 15 | - |  |  |  |
| Age at examination (years) | 32.2 (26,51) | 49.5 (30,65) | 0.054 |  |  |  |
| Sex %M | 60 | 37.5 | 0.38 |  |  |  |
| MRC Muscle grade (max 60) | 60 (60,60) | 51.9 (26,60) | 1.50E-05 |  |  |  |
| D4Z4 repeat length | - | 6.6 (3,9) | - |  |  |  |
| Ricci Score (max 10) | - | 4.4 (2,8) | - |  |  |  |
| Lamperti Score (max 15) | - | 5.1 (1,11) | - |  |  |  |
| Disease Duration | - | 22.5 (5,52) | - |  |  |  |
| Lower Limb fat fraction (%) | - | 15.3 (3.3,31.2) | - |  |  |  |
| MVC of tibialis anterior (N) | - | 66.8 (9.2,162.0) | - |  |  |  |

The FSHD severity indicators encompassed 3 clinical assessments (Ricci, Lamperti and MRC sum scores), an objective assessment of muscle strength (MVC of TA) and 3 variables shown to correlate with clinical severity (LLFF, disease duration, D4Z4 repeat length). Cross validation of these indicators had not previously been performed in FSHD.

All three clinical assessments correlated strongly with one another (Ricci vs Lamperti score: Pearson’s *r*=0.89, *p*=1.4×10^−9^; Ricci vs MRC sum score Pearson’s *r*=-0.72, *p*=2.9×10^−5^; Lamperti vs MRC sum score: Pearson’s *r*=-0.73, *p*=2.3×10^−5^, **Figure 1**). The three clinical severity scores also correlated with the objective measure of muscle strength *i*.*e*. MVC of TA (Ricci score vs MVC of TA: Pearson’s *r*=-0.53, *p*=0.011; Lamperti score vs MVC of TA: Pearson’s *r*=-0.52, *p*=0.013; MRC sum score vs MVC of TA: Pearson’s *r*=0.62, *p*=0.0022, **Figure 1**), as well as with LLFF (Ricci score vs LLFF: Pearson’s *r*=0.61, *p*=0.0013; Lamperti score vs LLFF: Pearson’s *r*=0.60, *p*=0.0016; MRC sum score vs LLFF: Pearson’s *r*=0.63, *p*=7.4×10^−4^, **Figure 1**). D4Z4 repeat length was inversely associated with LLFF (Pearson’s *r*=-0.57, *p*=0.0037, **Figure 1**), consistent with known association between shorter D4Z4 repeat lengths and more severe pathology^30^, although not with Ricci, Lamperti, MRC sum score nor MVC of TA. Self-reported FSHD disease duration did not correlate with any other markers of clinical severity (**Figure 1**).

**Figure 1:**
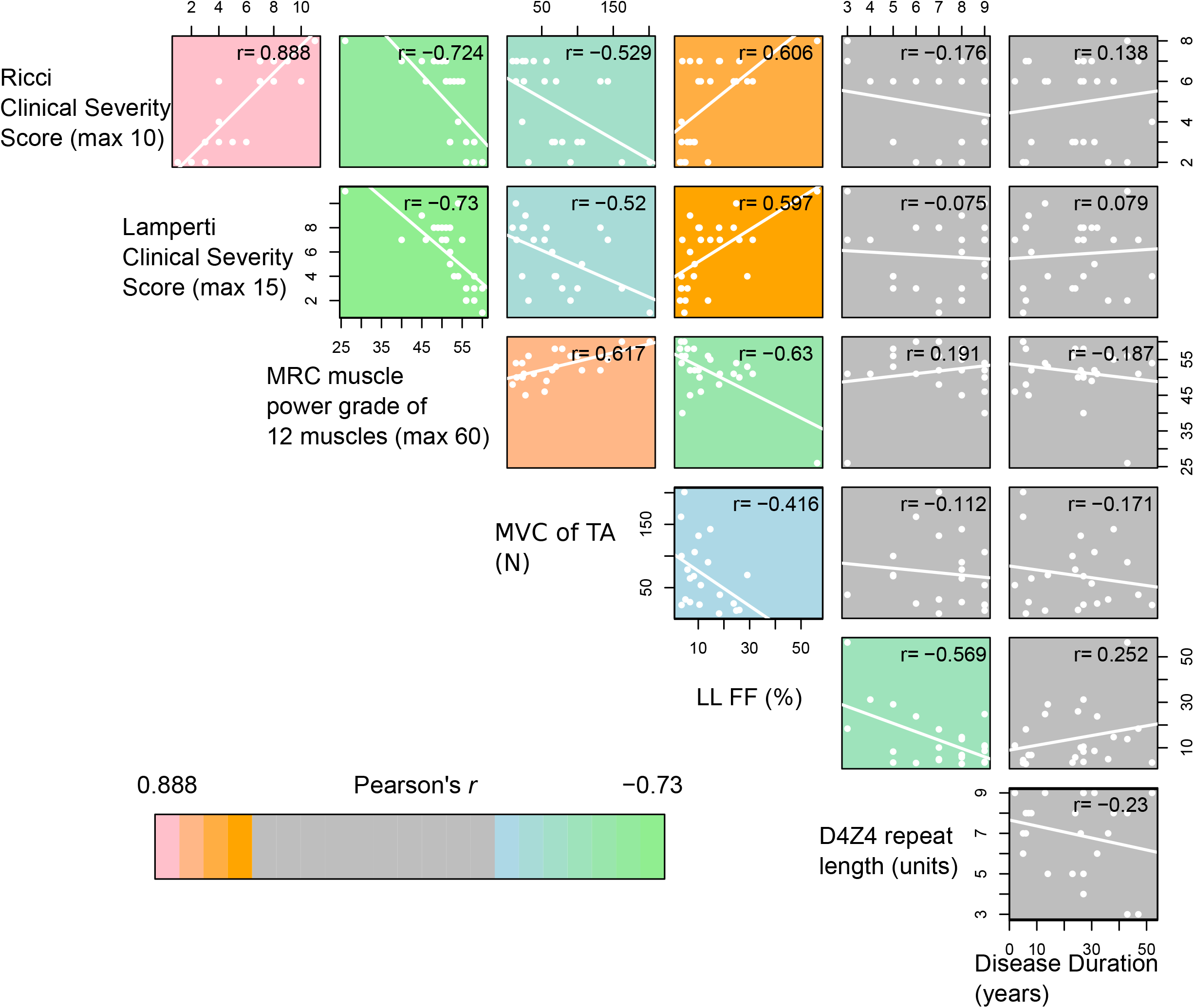
Correlation between measures of FSHD clinical severity. Scatterplots display comparisons of FSHD clinical severity scores: Ricci, Lamperti and MRC sum scores, MVC of TA, LLFF, D4Z4 repeat length (in FSHD1 individuals) and Disease duration. Comparisons with non-significant associations are coloured grey, while positive associations are coloured pink to orange and negative associations green to blue in order of significance (assessed at the 5% level). Pearson’s *r* for each comparison is displayed.

Multivariate linear regression showed Ricci and Lamperti scores were both higher in older male patients (Ricci score: age at examination *p*=0.012, sex *p*=0.033; Lamperti score, age at examination *p*=0.047, sex *p*=0.038). The MRC sum score was also significantly lower in male patients (*p*=0.042), but was not associated with age at examination. No other clinical variables demonstrated association between age or sex.

### PAX7 target gene repression discriminates control, inflamed (TIRM+) and non-inflamed (TIRM-) FSHD muscles

Muscle biopsies were obtained from 35/49 individuals for RNA-sequencing. 23 FSHD1 patients and 1 FSHD2 patient underwent MRI guided muscle biopsies with samples taken from TIRM-(23/24 vastus lateralis) and TIRM+ muscle (15/24 gastrocnemius medialis) from each patient (Table S1). 11 control individuals also underwent muscle biopsy of vastus lateralis. FSHD patients who underwent muscle biopsy were older than controls and showed a male sex bias (Table 1).

Five known FSHD biomarkers^9–11,17^ were considered in our muscle biopsy samples. Three biomarkers are based on DUX4 target gene expression: Choi et al., a set of 212 “early” DUX4 target genes^10 31^; Geng et al., a set of 165 “late” DUX4 target genes^10 32^ and Yao et al., a set of 114 “late” DUX4 target genes^33,34^. The fourth FSHD biomarker is based on PAX7 target gene repression and derived from 311 upregulated and 290 downregulated PAX7 target genes^10^. The final biomarker was our Lymphoblast score, derived from 237 genes found to be upregulated in FSHD lymphoblastoid cell lines compared to controls^11^.

As expected, PAX7 target gene repression proved a clear biomarker of FSHD status, discriminating FSHD and control muscle biopsies (Multivariate regression on all samples: PAX7 score vs Ctrl/TIRM-/TIRM+ status adjusting for age and sex *p*=0.0013).Our PAX7 biomarker also discriminated matched TIRM- and TIRM+ FSHD samples (Multivariate regression on FSHD samples: PAX7 score vs TIRM status adjusting for age, sex and patient *p*=0.013) (Figure 2A). The 3 DUX4 target gene biomarkers all discriminated FSHD from controls, however, none discriminated between matched TIRM- and TIRM+ FSHD samples on paired analysis at the 5% significance level (Figure 2B-D).

**Figure 2:**
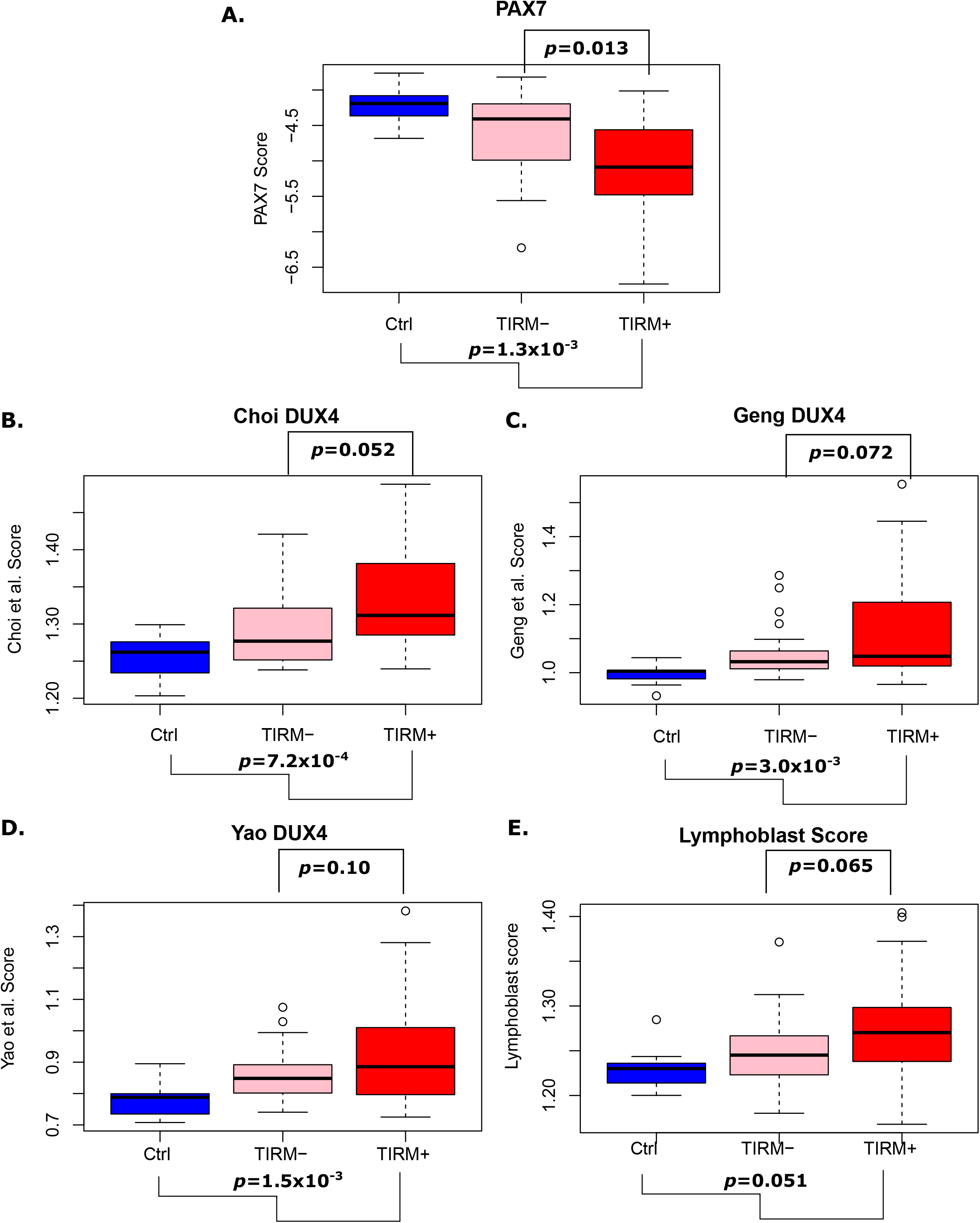
Analysis of FSHD Biomarkers in inflamed (TIRM+), non-inflamed (TIRM-) and control muscle and PBMCs. (A-E) Boxplots demonstrate (A) the PAX7 target gene score, (B) Choi et al. early DUX4 target genes, (C) Geng et al. and (D) Yao et al. late DUX4 target genes and (E) the Lymphoblast score in muscle biopsies from 11 control individuals and 24 FSHD TIRM- and TIRM+ samples. Box represents the IQR, with median indicated by a line. Whiskers denote min [1.5□IQR, max (observed value)]. Multivariate regression *p*-values adjusting for age and sex are shown for each biomarker as a correlate of control vs TIRM-/TIRM+ status below each plot and separately also adjusting for matched paired TIRM- and TIRM+ samples above each plot.

Our Lymphoblast score correlates with the degree of inflammation in FSHD muscle biopsies^11^. Here, the Lymphoblast score was not a significant biomarker of FSHD compared to control, nor was it able to discriminate between matched TIRM- and TIRM+ FSHD samples on paired analysis (Figure 2E).

A recent analysis investigated the PAX7 and DUX4 target gene biomarker sets in FSHD PBMCs via the *globaltest* package^23^. Unlike our approach, *globaltest* does not give a single sample score for use as a biomarker, but determines the association between a set of genes and given phenotype. We assessed the gene sets on which the 5 biomarkers are based using *globaltest* methodology, which did not change our findings for the PAX7, Geng et al and Yao et al DUX4 target genes. However, the Choi et al., early DUX4 target genes were also significantly associated with TIRM status among matched pairs of FSHD samples (adjusting for age and sex), and the genes comprising the Lymphoblast score were significantly associated with FSHD vs Control status as well as TIRM status in matched samples (Table S2). This indicates that our biomarker score construction is suboptimal for the Choi et al., DUX4 target gene signature and the Lymphoblast signature.

TIRM+ and TIRM-muscle biopsies were not all sampled from the same anatomical muscle. To determine the impact of anatomical muscle on the FSHD biomarkers we performed a regression analysis separately on TIRM- and TIRM+ FSHD biopsies, modelling each biomarker as a function of anatomical muscle, age and sex. We found no association between PAX7 target gene repression or the Lymphoblast score and anatomical muscle across TIRM+ and TIRM-samples and no association between the Choi et al., DUX4 target score and anatomical muscle across TIRM-samples. However, the Geng et al., and Yao et al., DUX4 target scores both showed elevated levels in the single *vastus intermedius* TIRM-sample compared to the 23 *vastus lateralis* TIRM-samples. The Yao et al., DUX4 target score also demonstrated elevated levels on the single *soleus* and *sartorius* TIRM+ samples compared to the other TIRM+ samples, while the Geng et al DUX4 score was elevated on the *sartorius* TIRM+ sample and the Choi et al., DUX4 score on the *soleus* TIRM+ sample, compared to other TIRM+ muscles. To limit impact of anatomical muscle on biomarker discrimination, data was re-analysed. The TIRM-*vastus intermedius* sample was excluded from comparison involving the Yao et al. and Geng et al. scores, the *satorius* TIRM+ sample was excluded from comparisons involving the Yao et al. and Geng et al. scores and the *soleus* sample was excluded from comparisons involving the Yao et al. and Choi et al. scores. Despite this, the significance of all results remained unchanged.

### PAX7 target gene repression correlates with Ricci and Lamperti clinical severity scores

We next considered association between FSHD biomarkers and clinical severity variables. Across TIRM-FSHD muscle, PAX7 target gene repression correlated with Ricci and Lamperti clinical severity scores (Ricci: Pearson’s *r*=-0.42, *p*=0.043; Lamperti: Pearson’s *r*=-0.41, *p*=0.045, **Figure 3A and B**). The Yao et al., late DUX4 target genes correlated with LLFF in TIRM-FSHD samples (Pearson’s *r*=0.51, *p*=0.012, **Figure 3C**).

**Figure 3:**
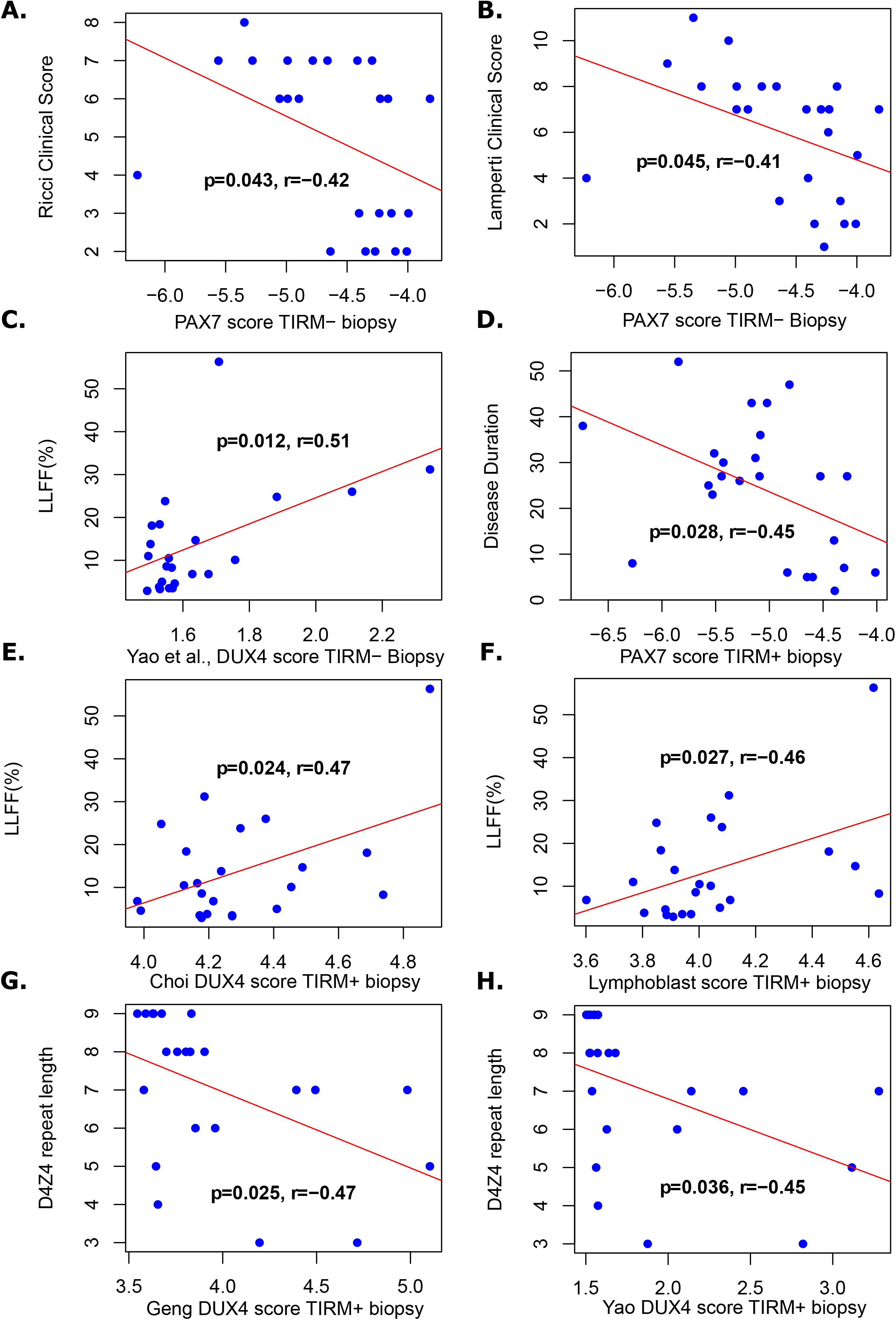
Association between FSHD biomarkers and clinical severity variables in muscle biopsies. Scatter plots show (A) Ricci and (B) Lamperti clinical severity scores against PAX7 target gene expression in TIRM-FSHD muscle biopsies from 24 patients. (C) LLFF is plotted against Yao et al. late DUX4 target biomarker in the same 24 TIRM-FSHD muscle biopsies. (D) Disease duration against PAX7 target gene expression in TIRM+ FSHD muscle biopsies from 24 patients. LLFF is plotted against (E) Choi et al., early DUX4 target biomarker and (F) the Lymphoblast score, in 24 TIRM+ FSHD muscle biopsies. D4Z4 repeat length is plotted against (G) Geng et al. and (H) Yao et al. late DUX4 target gene biomarkers in 23 FSHD1 TIRM+ muscle biopsies. Pearson’s *r* and associated *p*-value is given alongside a line of best fit.

Across TIRM+ FSHD muscle, PAX7 target gene repression correlated with disease duration (Pearson’s *r*=-0.45, *p*=0.028, **Figure 3D**), consistent with our demonstration that PAX7 target genes are progressively repressed in biopsies from the same patients taken one year apart^17^. Both Choi et al., early DUX4 target genes and the Lymphoblast score correlated with LLFF (Choi et al., Pearson’s *r*=0.47, *p*=0.024, **Figure 3E**; Lymphoblast score, Pearson’s *r*=0.46, *p*=0.027, **Figure 3F**). Lastly, late DUX4 target genes of Geng et al. and Yao et al. inversely correlated with D4Z4 repeat length (Geng et al., Pearson’s *r*=-0.47, *p*=0.025, **Figure 3G**; Yao et al., Pearson’s *r*=-0.45, *p*=0.036, **Figure 3H**).

### Generating an effective FSHD blood biomarker

PBMCs were isolated from 14 FSHD1 and 1 FSHD2 patients (12 FSHD1 and the FSHD2 also underwent muscle biopsies) and 14 control individuals (2 also underwent muscle biopsy). None of the five biomarkers discriminated FSHD and control PBMCs after adjustment for age and sex either via multivariate regression or *globaltest* (**Table S2**).

We considered each of our 5 biomarkers in the 13 FSHD PBMC samples with corresponding muscle biopsies, and investigated whether the level of the biomarkers correlated between muscle and blood. We found 2 significant positive correlations: between PAX7 target gene repression in PBMCs and TIRM-FSHD muscle biopsies (*p*=0.002, Pearson’s *r*=0.76, **Figure 4A**), and between the Lymphoblast score in PBMCs and TIRM+ FSHD muscle biopsies (*p*=0.04, Pearson’s *r*=0.57, **Figure 4B**). None of the DUX4 target gene expression biomarkers correlated in levels between muscle and PBMCs.

**Figure 4:**
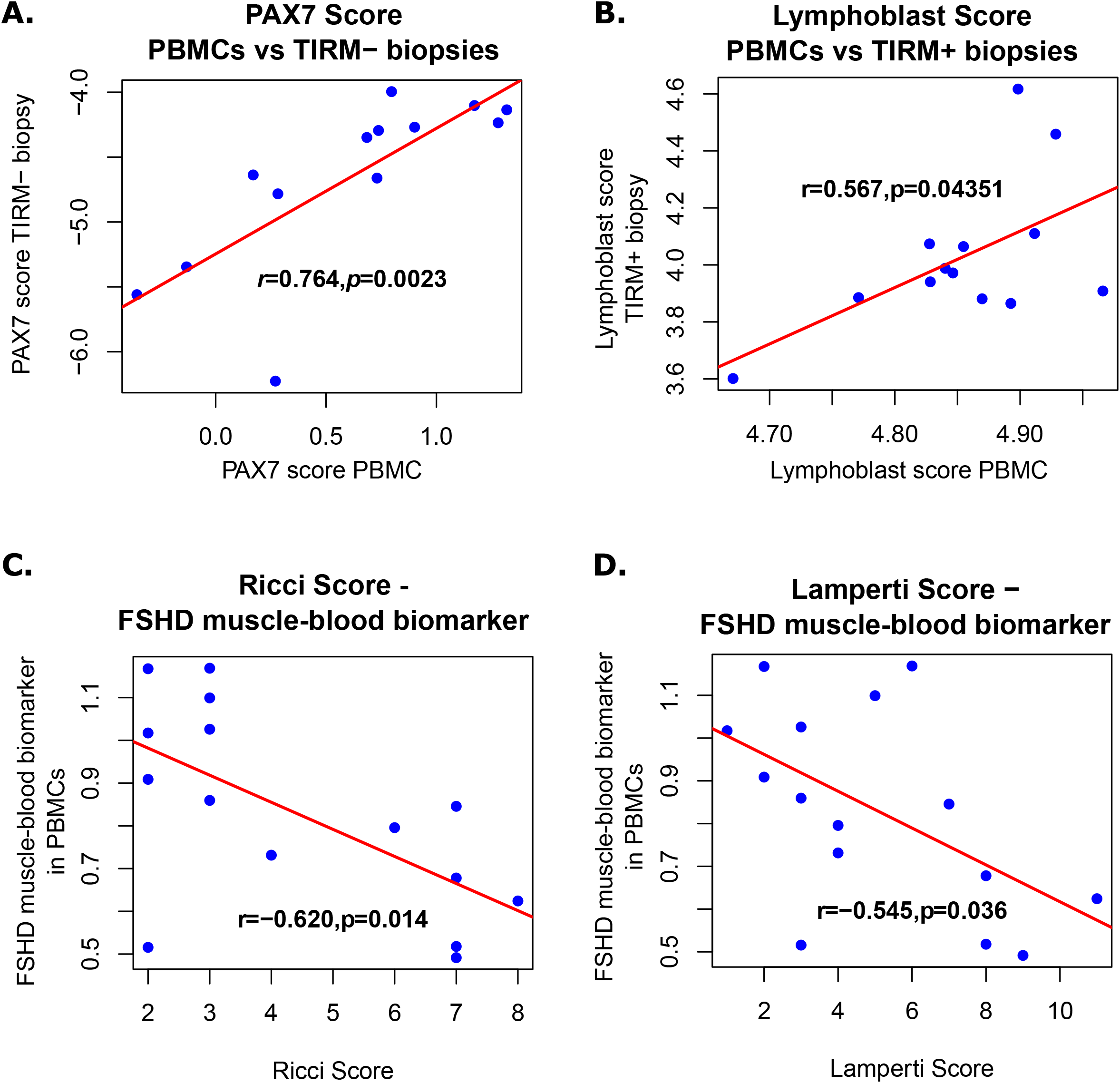
Association between PAX7 and Lymphoblast score across muscle and blood samples and association between FSHD muscle-blood biomarker in PBMCs and clinical severity measures. Scatter plots show (A) PAX7 target gene biomarker levels in 13 TIRM-FSHD muscle biopsies against levels in 13 matched PBMCs and (B) the Lymphoblast score levels in 13 TIRM+ FSHD muscle biopsies against levels in 13 matched PBMCs. The 143 gene FSHD muscle-blood biomarker in PBMCs is plotted against (C) Ricci and (D) Lamperti clinical severity scores in 15 FSHD PBMCs. Pearson’s *r* and associated *p*-value is provided, alongside line of best fit.

The levels of PAX7 target genes and Lymphoblast score in PBMCs, however, did not associate with any clinical variables. We thus performed a refinement of these two signatures to reduce noise in the association between the biomarker level in muscle and clinical variables.

For each of the 237 genes of the Lymphoblast score in TRIM- and TRIM+ muscle, we identified which had expression levels significantly correlated with the associated clinical variable of LLFF and identified 44/237 genes. A refined Lymphoblast score biomarker was set as the mean expression of the 44 genes in a given sample. The refined Lymphoblast score had no association with any clinical variables in PBMCs.

We next considered the 601 genes comprising the PAX7 score (311 up-regulated, and 290 down-regulated, PAX7 target genes). For each of the PAX7 target genes in TIRM- and TIRM+ muscle we identified which had expression levels significantly correlated with the associated clinical variables of the Ricci or Lamperti severity scores or disease duration. Using this analysis we identified 143 genes (64/311 up-regulated, 79/290 down-regulated, **Table S3**).

### The FSHD muscle-blood biomarker correlates with clinical severity in FSHD muscle and PBMCs

A refined PAX7 target gene biomarker was defined as the *t*-statistic evaluating the difference between expression of the 64 up-regulated PAX7 target genes and the 79 down-regulated PAX7 target genes associated with clinical severity in muscle, in a given sample. This refined PAX7 target gene biomarker showed significant association between its level in PBMCs and the Ricci or Lamperti clinical severity scores, independent of age or sex (Ricci score: *p*=0.014, *r*= -0.62; Lamperti score: *p*=0.036, *r*= -0.545, *Figure 4C and D*). PAX7 does not have a defined role in blood cells and the 143 genes identified are not necessarily direct targets of PAX7 and may be regulated in multiple ways. We therefore named our refined PAX7 biomarker the *FSHD muscle-blood biomarker*.

As per the full PAX7 score, our 143 gene FSHD muscle-blood biomarker can discriminate control, TIRM+ and TIRM-samples, (**Figure 5A**) and correlates in levels between PBMCs and TIRM-muscle biopsies (**Figure 5B**). Importantly, using the longitudinal, transcriptomic muscle biopsy dataset published by Wong et al. 2019^14^ the FSHD muscle-blood biomarker discriminates year 1 from year 2 FSHD muscle biopsy samples (paired Wilcoxon *p*=0.03, **Figure 5C**), as does the full PAX7 score. Finally, the FSHD muscle-blood biomarker discriminates FSHD from control samples on meta-analysis and on 6/7 independent FSHD muscle biopsy transcriptomic datasets (Fisher’s combined *p*=3.4×10^−10^, **Figure 5D**), a performance comparable to the full PAX7 score (significant on all datasets).

**Figure 5:**
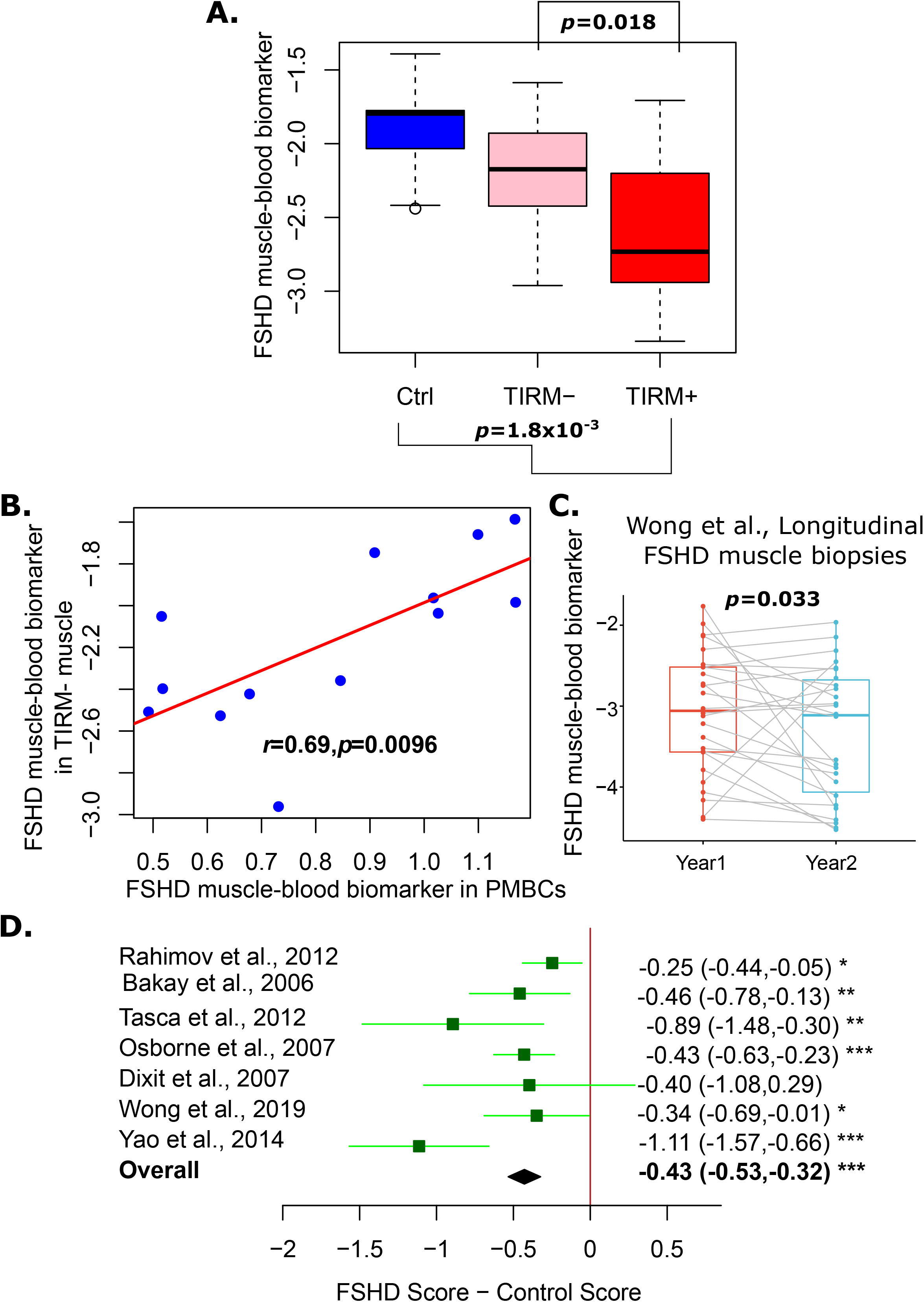
Validation of the 143 gene FSHD muscle-blood biomarker. (A) Boxplot demonstrates the 143 gene FSHD muscle-blood biomarker in muscle biopsies from 11 control individuals and 24 FSHD TIRM- and TIRM+ samples. Box represents the IQR, with median indicated by a line. Whiskers denote min [1.5□IQR, max (observed value)]. Multivariate regression *p*-values adjusting for age and sex are shown for the biomarker as a correlate of control vs TIRM-vs TIRM+ status below the plot and separately also adjusting for matched paired TIRM- and TIRM+ samples above the plot. (B) Scatterplot displays the FSHD muscle-blood biomarker in 13 TIRM-FSHD muscle biopsies against levels in 13 matched PBMCs. Pearson’s *r* and associated *p*-value is given alongside a line of best fit. (C) Boxplot demonstrates the refined 143 gene PAX7 score in 26 matched year 1 and year 2 FSHD muscle biopsy samples described by Wong et al.,^14^. Lines demonstrate connections between paired samples and a paired Wilcoxon test *p*-value is displayed. Box represents the IQR, with median indicated by a line. Whiskers denote min [1.5□IQR, max (observed value)]. (D) Forest plot displays the significance of the FSHD muscle-blood biomarker as a discriminator of FSHD muscle biopsies in seven independent microarray or RNA-seq datasets (130 FSHD, 98 control). Boxes denote mean difference in the FSHD muscle-blood biomarker between FSHD and control muscle biopsies and whiskers denote 95% confidence interval. A vertical line denotes a score difference of 0 and datasets where the whiskers cross this line have not attained significance at *p*<0.05 (as assessed by Wilcoxon test). Numerical values for mean score difference and confidence interval are displayed for each dataset to the right of the plot with significance denoted by asterisks where * denotes *p*<0.05, ** denotes *p*<0.01 and *** denotes *p*<0.001. Overall estimate is displayed as a diamond and was computed using a random effects model with significance assessed via Fisher’s combined test.

GSEA via Fisher’s exact test on genes comprising the FSHD muscle-blood biomarker was performed. Genes suppressed in muscle and blood of patients with severe disease showed enrichment for pathways previously implicated in FSHD: including targets of the DREAM complex^35^, neuronal gene sets^36^, Wnt signalling^37^, hypoxia response^10^ and hormone response and regulation^38,39^, including 17β-hydroxysteroid dehydrogenase 8 targets, and estradiol response^40^ (**Table S4**). Genes up-regulated in muscle and blood of patients with severe disease showed enrichment for epithelial to mesenchymal transition^41^, Wnt signalling^37^, TGF-β signalling^42^ and vasculature development^43^ (**Table S4**).

### The FSHD muscle-blood biomarker validates as a blood biomarker of clinical severity in an independent data set

To validate our FSHD muscle-blood biomarker as a circulating measure of clinical severity in FSHD we considered the recent data set of Signorelli et al., 2020^23^ describing RNA-sequencing of PAXgene blood from 54 FSHD patients and 29 controls. The dataset comprises 2 independent cohorts processed separately. The Nijmegen cohort comprises 39 FSHD and 11 control samples. The Newcastle cohort comprises 15 FSHD and 18 control samples. Clinical annotations for each sample included patient age, sex and the Lamperti clinical severity score. Raw read counts were obtained from the authors and data was normalised separately in each cohort, values for the FSHD muscle-blood biomarker were computed for each sample and *z*-normalised within cohort.

The FSHD blood-muscle biomarker was not a significant indicator of FSHD status on our dataset of 15 FSHD and 14 control PBMCs, after adjusting for age and sex (Multivariate regression *p*=0.2), but demonstrated lower average values on FSHD samples (FSHD mean: 0.83, range: (0.49,1.16), control mean: 0.93, range: (0.67,1.2)). To determine association of the FSHD muscle-blood biomarker with FSHD status in the larger dataset of Signorelli et al., multivariate regression analysis was performed modelling the *z*-normalised FSHD muscle-blood biomarker values as a function of FSHD status, age, sex and cohort. The FSHD muscle-blood biomarker was significantly lower on FSHD blood samples versus controls, independently of age, sex and cohort (*p*=0.002, **Figure 6A**).

**Figure 6:**
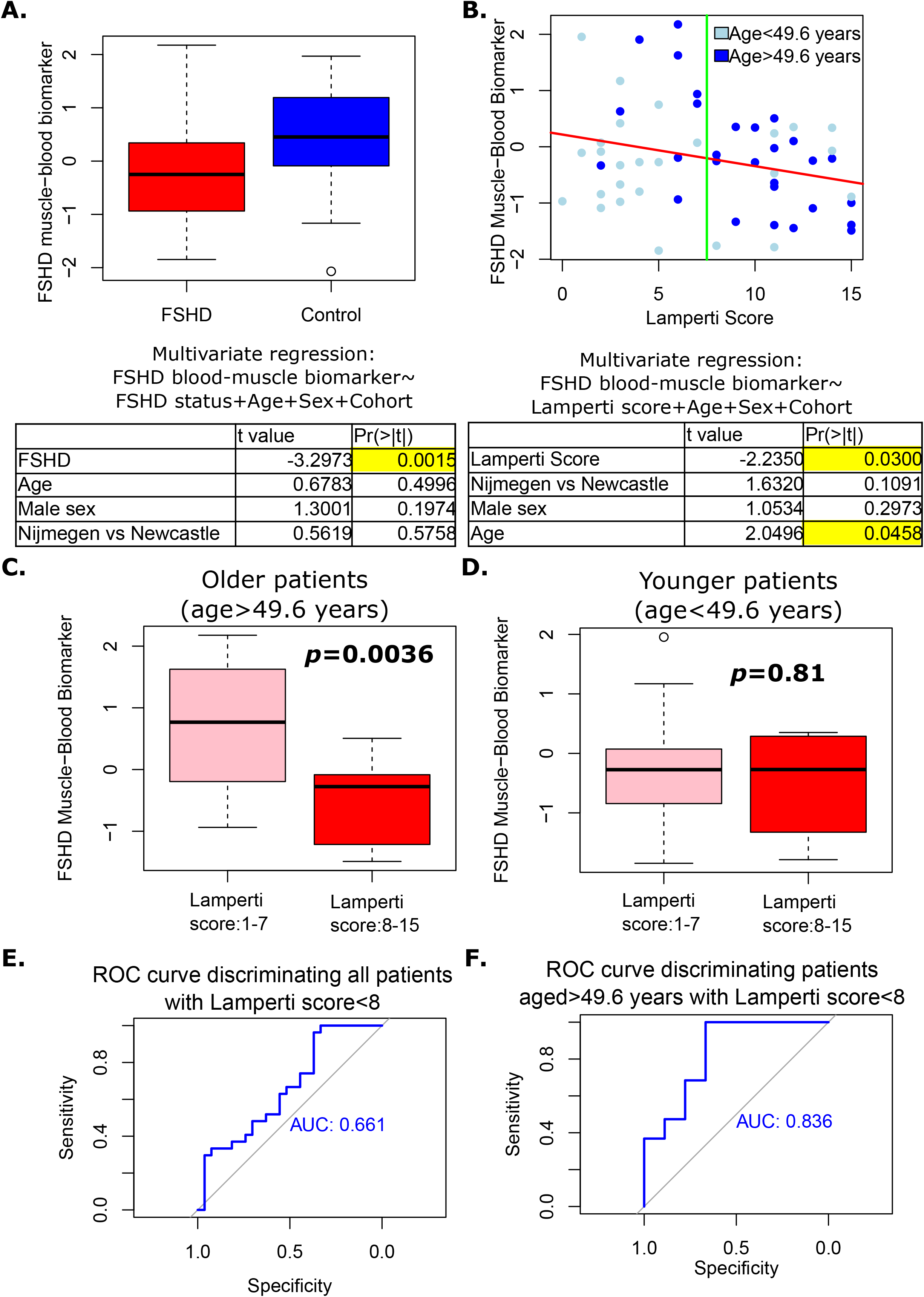
Investigation of the FSHD muscle-blood biomarker in 83 FSHD and control peripheral blood samples. (A) Boxplot demonstrates the 143 gene FSHD muscle-blood biomarker, z-normalised within cohort in peripheral blood RNA-seq samples from 54 FSHD patients and 29 controls described by Signorelli et al., 2020 ^23^. Box represents the IQR, with the median indicated by a line. Whiskers denote min [1.5□IQR, max (observed value)]. Below the plot a table summarises the results of multivariate linear regression, modelling the FSHD muscle-blood biomarker across 83 samples as a function of FSHD status, age, sex and cohort. (B) Scatter plot displays the FSHD muscle-blood biomarker, *z*-normalised within cohort in peripheral blood RNA-seq samples of 54 FSHD patients against Lamperti score. Light blue points denote patients below, and dark blue for patients above, the mean cohort age of 49.6 years. Red line of best fit is presented and a green line corresponds to Lamperti score=7.5, separating patients with mild and severe FSHD. The table summarises the results of multivariate linear regression, modelling the FSHD muscle-blood biomarker across 54 FSHD samples as a function of Lamperti score, age, sex and cohort. Boxplots display the FSHD muscle-blood biomarker in (C) 18 FSHD patients with Lamperti scores 1-7 and 8 FSHD patients with Lamperti scores 8-15 aged<49.6 and (D) 9 FSHD patients with Lamperti scores 1-7 and 19 FSHD patients with Lamperti scores 8-15 aged>49.6. Box represents the IQR, with median indicated by a line. Whiskers denote min [1.5□IQR, max (observed value)]. Multivariate regression analysis of FSHD muscle-blood biomarker as a function of FSHD severity (Lamperti score 1-7 vs 8-15), age, sex and cohort was performed separately in younger and older groups, *p*-values for the severity coefficient are displayed on each plot. ROC curves and associated AUC values for the FSHD muscle-blood biomarker as a marker of FSHD severity (Lamperti score 1-7 vs 8-15) are displayed for (E) all 54 FSHD samples and (F) 28 FSHD patients aged >49.6 years.

The FSHD muscle-blood biomarker significantly correlated with the Lamperti score in our dataset of 15 FSHD PBMCs. To determine association of the FSHD muscle-blood biomarker and Lamperti score in the larger blood sample dataset of Signorelli et al., multivariate regression analysis was performed modelling the *z*-normalised FSHD muscle-blood biomarker values as a function of Lamperti score, age, sex and cohort. The FSHD muscle-blood biomarker significantly negatively correlated with the Lamperti score independently of age, sex and cohort (*p*=0.030, **Figure 6B**). Confirming our biomarker as a measure of clinical severity in blood samples.

To ascertain whether this association is attributable to chance, we performed resampling, selecting 1000 independent, random, non-overlapping sets of 143 genes, divided into 64 up-regulated and 79 down-regulated targets. For each random gene-set we performed multivariate analysis as above to determine association with FSHD status and severity in the Signorelli et al. dataset. Only 32/1000 random signatures could discriminate FSHD from control samples as well as the true FSHD muscle-blood biomarker (*p*=0.032), and only 6/1000 showed as strong an association with the Lamperti score (*p*=0.006).

Curiously, the FSHD muscle-blood biomarker positively correlated with patient age, independently of Lamperti score, sex and cohort (*p*=0.046, **Figure 6B**). This is unexpected as Lamperti score and patient age are positively correlated in the Signorelli et al. dataset (Pearson’s *r*=0.35, *p*=0.009). It would be anticipated that the FSHD muscle-blood biomarker would have unidirectional association with both variables. This indicates that the FSHD muscle-blood biomarker has different associations with the Lamperti score in older versus younger patients. To investigate, we split the FSHD patients in the Signorelli et al., dataset into 2 groups based on mean age of FSHD patients in the cohort (49.6 years): an older group (28 patients aged>49.6 years) and a younger group (26 patients aged<49.6 years). The younger and older groups showed no significant difference in sex distribution (*p*=0.74) or Nijmegen/Newcastle cohort (*p*=0.64). As expected, the older group displayed significantly higher Lamperti scores (*p*=0.002, younger group mean: 5.8 range: 0-15, older group mean: 9.4, range:2-15, **Figure 6B**). Association between the FSHD blood-muscle biomarker and the Lamperti score is driven by the older patient cohort, with the score robustly able to discriminate older patients with mild FSHD (Lamperti score: 1-7) from those with more severe FSHD (Lamperti score: 8-15) (*p*=0.004, **Figure 6C**), but showing no discrimination of mild versus severe in younger patients (*p*=0.81, **Figure 6D**). As a binary classifier of mild versus severe FSHD, the FSHD muscle-blood biomarker obtains modest accuracy on the full Signorelli et al., dataset (AUC=0.661, **Figure 6E**), but robust classification on the older FSHD patients (AUC=0.836, **Figure 6F**).

## Discussion

We investigated transcriptomic biomarkers in RNA-sequencing data from paired non-inflamed (TIRM-) and inflamed (TIRM+) muscle biopsies and PBMCs from clinically characterised FSHD patients, alongside matched controls. Our PAX7 target gene repression biomarker^10^ discriminates control, inflamed and non-inflamed FSHD muscle, while the discriminatory power of DUX4 target gene signatures is limited to distinguishing control from FSHD muscle. We also developed a new 143 gene FSHD muscle-blood biomarker that shows levels correlated between paired TIRM-muscle and PBMCs. Importantly, the level of our FSHD muscle-blood biomarker in both TIRM-muscle and PBMCs correlates with Ricci and Lamperti measures of clinical severity. By analysing a published dataset describing RNA-sequencing of peripheral blood for 54 clinically characterised FSHD patients and 29 matched controls, we validated our FSHD muscle-blood biomarker as a circulating measure of clinical severity, with particular strength in older patients.

Our FSHD muscle-blood biomarker is the first circulating indicator of FSHD clinical severity with direct correlation to transcriptomic changes in patient muscle, valid on a cohort level. The signature could be developed into a minimally-invasive tool for routine use in clinic and monitoring patients in trials. The 143 genes comprising the FSHD muscle-blood biomarker represent an important discovery set for FSHD pathomechanisms and therapeutic targets, valid in both muscle and blood.

FSHD displays both inter-patient and intra-patient heterogeneity, with the degree of muscle weakness and affected muscle distribution varying dramatically. This inhibits development of reliable biomarkers based on single muscle biopsies from patients. Our well-controlled dataset takes two biopsies from each clinically characterised FSHD patient alongside paired peripheral blood, allowing adjustment for heterogeneity and identification of a transcriptomic biomarker valid in multiple tissue types.

Importantly, our FSHD muscle-blood biomarker correlated with robust clinical assessments. Comparative analysis of clinical severity measures for FSHD demonstrated strong concordance between the Ricci and Lamperti clinical severity scores, which in turn correlate with objective measures of function in FSHD patients including MVC of TA and LLFF on MRI, confirming the clinical assessments as robust characterisations of FSHD severity.

The dataset of Signorelli et al., 2020^23^ was used as a validation set for our biomarkers and we confirmed findings that the full PAX7 target gene score and DUX4 target gene sets are not FSHD blood biomarkers. Signorelli et al., reported no genes with expression levels associated with FSHD status or clinical severity in their discovery analysis when investigating the Nijmegen and Newcastle cohorts separately. In our validation approach to this data, we compute our novel FSHD muscle-blood biomarker separately in each cohort, and integrate the independent datasets, facilitating a larger sample size to validate our biomarker as a negative correlate of clinical severity.

We found that the FSHD muscle-blood biomarker has a stronger association with clinical severity in older patients. This may be attributed to several factors, but the FSHD muscle-blood biomarker could have predictive value on FSHD disease progression. If true, in older patients who have reached their potential disease severity, clinical scores and biomarker values may be expected to match closely, while in younger patients who are early in the disease process, low biomarker values may imply a more severe disease course progression. There are no patients (young or old) with high biomarker values (normalised biomarker level>0.567) and severe FSHD (Lamperti scores>7), implying no patients have been ‘predicted’ a mild score and achieved a severe score. Moreover patients with low biomarker values and mild FSHD (Lamperi scores<7) are younger (83% below mean cohort age), suggesting they may still attain a ‘predicted’ severe phenotype with age. Conversely those with low biomarker values and severe FSHD tend to be older (70% above mean cohort age), suggesting that a ‘predicted’ severe phenotype has been achieved by greater age. Validation of the FSHD muscle-blood biomarker as a predictor of severity will require longitudinal studies.

Recently, van den Heuvel et al., 2022^12^ performed RNA-sequencing of FSHD and control muscle biopsies from different muscle groups. The authors identified non-overlapping roles for PAX7 and DUX4 target genes, with both biomarkers independently associated with fatty replacement of muscle but only DUX4 target gene expression associated with inflammation^12^, consistent with our prior findings^9,11^. This emphasises importance of studying biomarkers in inflamed and non-inflamed muscle separately. As our study profiles paired TIRM+/TIRM-muscle biopsies from the same FSHD patients, we account for inter-patient heterogeneity in FSHD, allowing deeper analysis of biomarker associations. PAX7 target genes associate with FSHD muscle regardless of inflammatory state, and can discriminate TIRM+ FSHD muscle from paired TIRM-. By contrast, DUX4 target genes have discriminatory power limited to separating FSHD muscle from control. Van den Heuvel et al., highlighted importance of minimally invasive biomarkers for FSHD severity, focusing on use of MRI^12^. We propose a complementary approach, based on gene expression in minimally invasive blood samples.

We found no correlation between DUX4 target gene expression in FSHD muscle and paired PBMCs. Our Lymphoblast score comprises genes up-regulated in FSHD lymphoblastoid cell lines compared to matched controls^11^ and here demonstrated significant correlation between TIRM+ FSHD muscle and PBMCs, but not TIRM-biopsies. This is consistent with our previous demonstration that the Lymphoblast score associates with the degree of histological inflammation in FSHD muscle biopsies^11^. However, the Lymphoblast score could not be further refined into a biomarker of FSHD clinical severity valid in PBMCs, though this may be related to construction of the Lymphoblast score, which was shown suboptimal by *globaltest*. Curiously, our PAX7 target gene repression score showed strong correlation in levels between TIRM-FSHD muscle and PBMCs, but not TIRM+, supporting a role for PAX7 target genes specifically in non-inflamed muscle. Studies have proposed a role for PAX7 in development of the thymus^44,45^ and *DUX4* expression has been shown in healthy human thymus^46^, raising the possibility that the two proteins may interact in lymphoid progenitor tissue and possibly affect epigenetic programming, as has been proposed in muscle^15^. PAX7 is not known to be expressed in mature immune cells, although Pax5 is expressed in lymphoid precursors and required for B-cell differentiation^47^.

The PAX7 target gene biomarker was refined to the FSHD muscle-blood biomarker, a measure of clinical severity valid in TIRM-muscle and PBMCs. Genes comprising the PAX7 target gene biomarker are not necessarily direct PAX7 targets and likely regulated by multiple transcription factors/other events. GSEA of the 143 genes comprising the FSHD muscle-blood biomarker identified numerous pathways known to be dysregulated in FSHD, including vasculogenesis, and hypoxic response, indicating that misregulation of these processes in FSHD is not limited to muscle. TGF-β signalling associated with genes up-regulated in clinically severe FSHD and is well-studied in muscular dystrophy, particularly myostatin which limits muscle growth via mechanisms directly relevant to FSHD, including direct suppression of PAX7 and activation of p38^48^. Targeting TGF-β has been considered in FSHD via the small molecule inhibitor ACE-083. Phase II trials demonstrated improvement of primary end-point (muscle mass of TA and biceps brachii), failure to improve secondary functional outcomes led to abandonment of trials^42^.

A recent trial of p38 inhibitor losmapimod^18^ designed to inhibit DUX4 activity, failed to demonstrate improvement of primary end-point (activation of 4 DUX4 target genes), but did demonstrate improvement of secondary functional measures. It would be interesting to measure our FSHD muscle-blood biomarker in blood/muscle samples from patients from this trial.

In summary, we report the FSHD muscle-blood biomarker a measure of clinical severity valid in both muscle biopsies and peripheral blood cells. Validation in an independent cohort confirmed our circulating measure as a correlate of disease severity, with particular validity in older patients, raising possibility of predictive value. The FSHD muscle-blood biomarker may represent a powerful tool for disease monitoring in trials and routinely in clinical assessment.

## Methods

### Participants

FSHD patients (n=25 FSHD1 and 1 FSHD2) and unrelated healthy control individuals (n=23) were recruited from 2019 to 2021 at the neurology outpatient clinic, Radboud University Medical Center, Nijmegen, The Netherlands. Patient criteria: 1) genetically confirmed FSHD, 2) age ≥ 18 years, 3) absence of infectious and/or inflammatory comorbidities, 4) absence of autoimmune comorbidities, 5) absence of past/present malignancy, 6) no use of corticosteroids, statins or anti-inflammatory medication. Unrelated healthy individuals were: 1) age ≥ 18 years old, 2) negative personal/family medical history for neuromuscular disorders, 3) absence of muscle weakness during physical examination. Study was approved by the regional medical ethical committee approval (CMO Arnhem-Nijmegen). Subjects provided written informed consent.

### Clinical Assessments

Age, sex, height, weight, BMI, and MRC sum score were recorded. MRC sum score was calculated as the sum of MRC scores from six muscle regions on both sides (upper arm abductors, elbow flexors, wrist extensors, hip flexors, knee extensors and foot dorsal flexors). Total score ranges from 0 (complete paralysis) to 60 (normal muscle strength). Additional clinical features were assessed for FSHD patients including the Ricci score^27^, the Lamperti score^29^ and maximum voluntary contraction (MVC) of tibialis anterior. Ricci score^27^ is a 0-10 severity scale that assumes a descending spread of symptoms from face and shoulders to pelvic and leg muscles typical of FSHD. The Lamperti score^29^ comprises a 0-15 severity scale evaluating degree of muscle weakness in 5 muscle regions separately. MVC of tibialis anterior was calculated as described by de Jong et al^49^. Finally, FSHD genetic characteristics (FSHD type and D4Z4 repeat length) and disease duration were obtained.

### PBMCs collection

PBMCs were isolated by dilution of fresh EDTA blood in sterile phosphate-buffered saline (PBS) and density centrifugation over Ficoll-Paque (GE healthcare, Zeist, The Netherlands) as described^50^.

### Magnetic resonance imaging

FSHD patients were examined on the same 3T MR system (Tim TRIO, Siemens, Erlangen, Germany) adapting the protocol of Mul et al.^51^. Briefly, Dixon and Turbo Inversion Recovery Magnitude (TIRM) sequences were made using a phased array birdcage coil around upper and lower legs. Dixon sequences quantified the degree of muscle fatty infiltration. Adjustments were made for the DIXON sequence: field of view (FOV) 435 mm, slice thickness 5 mm, gap 5 mm, repetition time (TR) 10 ms, time to echo (TE1 / TE2) 1,26 / 2,49 ms, number of slices per slab 72, flip angle (FA) 3 degree, base resolution 320. For TIRM sequence used to identify inflammation, an inversion time of 240 ms was selected to suppress the fat signal and parameters were: FOV 435 mm, slice thickness 5 mm, gap 10 mm, TR 4140 ms, TE 41 ms, number of slices per slab 28, FA 150 degree, base resolution 320.

### Muscle biopsy collection

We collected two paired muscle samples from each of 24 FSHD patients: one MRI-guided muscle biopsy targeting a TIRM negative (non-inflamed) muscle *vastus lateralis* (*vastus intermedius* from one patient) and a second sample from a leg muscle scored as TIRM hyperintense (presumed active inflammation). The area of biopsy was selected on: presence of TIRM hyperintensity, degree of fatty infiltration, amount and location of normal appearing muscle. Muscle biopsy site was marked on the skin with a fish oil marker, positioned on a reference line connecting the anterior superior iliac spine with the tibial tuberosity for upper leg, and the tibial tuberosity with the lateral malleolus for lower leg. Transversal 3D T1-weighted high resolution images (FOV 269 mm, slice thickness 1 mm, TR 759 ms, TE 2,61 ms, number of slices per slab 160, FA 13 degree, base resolution 384), TIRM (FOV 175 mm, slice thickness 4 mm, TR 4100 ms, TE 42 ms, number of slices per slab 23, FA 150 degree, base resolution 256), and DIXON images (FOV 500 mm, slice thickness 5 mm, TR 9,18 ms, TE1 / TE2 1,27 / 2,5 ms, number of slices per slab 52, FA 8 degree, base resolution 384), were made to confirm and determine exact area of biopsy. An experienced interventional radiologist determined needle trajectory and insertion site. Skin was cleaned with chlorhexidine in alcohol or other disinfectant. Biopsy site was infiltrated with 5 ml of 2% lidocaine taking care to inject skin and subcutaneous tissue, but not muscle. A 5 mm incision was made, and the skin layer penetrated with a scalpel blade. A coaxial needle containing a plastic introduction sheath and inner cutting stylet was introduced (ATEC, Hologic, Bedford, USA). The inner needle was then retracted and replaced by a blunt plastic obturator whereafter localizing images confirmed correct needle position, with trajectory adjusted towards target area as needed. After adjustment, fast verification images were made in at least two planes. The biopsy was taken using a MR compatible 9 gauge vacuum-assisted needle (ATEC, Hologic, Bedford, USA) and a verification image with the plastic obturator in situ confirmed biopsy site and evaluated possible complications. Finally, the sheath was removed and pressure applied over biopsy area to prevent bleeding. Steri-strips and bandage were applied to close the incision.

Bergström needle muscle biopsy from the *vastus lateralis* were collected from 11 control individuals as described^52^.

### MRI analysis

Dixon MRI sequences were analyzed using MATLAB (version R2020a, The Mathworks, Inc. Natick, Massachusetts, United States) and ImageJ software. A fat fraction (FF) map was calculated using MATLAB from the water and fat image of the Dixon sequence according to:

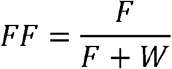

Muscle contours of 12 upper leg muscles (*sartorius, gracilis, vastus medialis, vastus lateralis, vastus intermedius, rectus femoris, biceps femoris caput brevis, biceps femoris caput longus, semitendinosus, semimembranosus, adductor magnus, adductor longus*) and 7 lower leg muscles (*tibialis anterior, extensor digitorum longus, peroneus, tibialis posterior, soleus, gastrocnemius medialis, and gastrocnemius lateralis*) muscles were manually outlined on the fat fraction map at specific regions as described^51^. Average fat fraction within one muscle contour 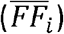 was calculated using ImageJ. Finally, average lower limb fat fraction (LLFF) weighted on the area (A) of each muscle was calculated according to:

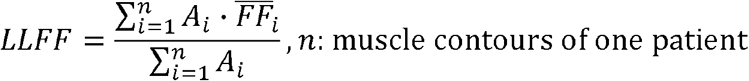

### Analysis of clinical variables

Clinical variables assessed included age at examination, sex and MRC muscle power grade. Variables were compared between FSHD and control individuals in 3 settings: entire cohort, subset of cohort in which muscle biopsies were performed and subset of cohort in which PBMCs were isolated. Age at examination and MRC sum score were compared in these 3 settings via Wilcoxon test, while sex was compared via logit regression, significance was assessed at the 5% level.

FSHD severity indicators: Ricci, Lamperti and MRC sum score, MVC of TA, LLFF, D4Z4 repeat length (FSHD1 patients) and Disease Duration, were compared across the 26 FSHD patients in the full cohort. Pearson correlation analysis was performed for each pairwise combination of the 7 variables, with significance assessed at *p*<0.05.

### Processing of RNA-sequencing data

RNA extraction from muscle biopsies, purification, quality control, library preparation and sequencing at 21.7-35.5 million reads/sample were performed by Genewiz (https://www.genewiz.com). RNA was extracted from PBMCs followed by globin depletion, quality control, library preparation and sequencing at 19.7-46.5 million reads/sample performed by Genewiz (https://www.genewiz.com).

Raw reads were trimmed using trim-galore, utilising cutadapt to remove Illumina Sequencing Adapters at the 3’ end. Additionally, 15 bases were trimmed from the 5’ end of the reads due to biased distributions. Reads were mapped to the human transcriptome using human genome sequence GRCh38 and v103 gene annotations downloaded from Ensembl, via the Salmon tool (v1.6.0) to correct for fragment GC content bias^53^, summarising data at the transcript level. For gene level analysis, read counts assigned to given transcripts were summed over in each sample. The resulting matrix of read counts was analysed using R.

### Computation and comparison of FSHD biomarkers

Computation of the three DUX4 expression biomarkers, PAX7 target gene repression biomarker and Lymphoblast score were as previously described^9–11,17^. Briefly, each DUX4 target gene expression score is computed for each sample as mean expression of genes found to be up-regulated by the studies of Yao et al.^34^ (114 genes), Geng et al.^32^ (165 genes) and Choi et al.^31^ (212 genes), normalised by their standard deviation within each sample. PAX7 target gene repression score for each sample was computed as the *t*-score from a test comparing up-regulated (311 genes) to down-regulated (290 genes) PAX7 target genes within each sample. We have published software for computation of each score from suitably normalized dataset^9^. The FSHD Lymphoblast score was computed in each sample as mean expression of 237 genes found up-regulated in FSHD Lymphoblastoid cell lines in our previous analysis^11^. Biomarker values in control samples (PBMCs, muscle biopsies) were compared with corresponding FSHD samples (PBMCs, TIRM- and TIRM+ muscle biopsies), via multivariate linear regression, modelling each biomarker as a function of age, sex and discrete variable encoding control (valued 0), TIRM-/PBMC (valued 1) and TIRM+ (valued 2) samples. Values in matched TIRM+ and TIRM-muscle biopsies were compared via a separate multivariate analysis including a patient covariate to adjust for paired samples. Significance was assessed at *p*<0.05.

Globaltest was implemented in R via the corresponding Bioconductor package^54^, analogous to the above multivariate analysis using 2 separate analyses. In the first the response vector was set via a discrete variable encoding control (valued 0), TIRM-/PBMC (valued 1) and TIRM+ (valued 2) and dependent variables were age and sex and genes comprising each biomarker. In the second analysis limited to FSHD samples the response vector described TIRM+ and TIRM-samples and dependent variables were age, sex, paired sample and genes comprising each biomarker. In both scenarios globaltest was implemented 5 times employing genes comprising the PAX7 score, the 3 sets of DUX4 target genes and the Lymphoblast score as covariates. Significance was assessed at the 5% level.

### Association between FSHD biomarkers and clinical severity variables and across tissues

For each of the 5 FSHD biomarkers: PAX7 target gene repression, Choi et al., Geng et al. and Yao et al., DUX4 target gene scores and the Lymphoblast score Pearson correlation analysis was performed separately in 24 TIRM- and 24 TIRM+ FSHD muscle biopsies to determine association between each of the FSHD clinical severity indicators: Ricci, Lamperti and MRC sum scores, MVC of TA, LLFF, D4Z4 repeat length (FSHD1 samples only) and disease duration. For each of the 5 biomarkers, Pearson correlation was also performed to determine association between biomarker levels in 13 TIRM- and TIRM+ FSHD muscle samples and corresponding levels in matched PBMCS. Significance was assessed at *p*<0.05.

### Constructing and correlating the refined target gene signatures

For each of the 601 genes comprising the full PAX7 target gene biomarker (311 up-regulated: 290 down-regulated) a Pearson’s correlation analysis was performed investigating association between expression of each gene in the 24 TIRM- and 24 TIRM+ FSHD samples separately and clinical variables: Ricci and Lamperti severity scores and disease duration – found to associate with the full PAX7 biomarker in muscle. We found 143/601 genes (64/311 up-regulated: 79/290 down-regulated) significantly correlated with one of the three clinical variables across either TIRM-or TIRM+ samples at *p*<0.05. These 143 genes were employed to construct the FSHD muscle-blood biomarker as the *t*-statistic evaluating the difference between expression of 64 up-regulated PAX7 targets and 79 down-regulated PAX7 targets.

For each of the 237 genes comprising the full Lymphoblast score a Pearson’s correlation analysis was performed investigating association between expression of each gene in the 24 TIRM- and 24 TIRM+ FSHD samples separately and LLFF – found to associate with the full Lymphoblast score in muscle. We found 44/237 genes significantly correlated with LLFF across either TIRM-or TIRM+ samples at the 5% level. The mean expression of these 44 genes constituted the refined Lymphoblast score.

Multivariate linear regression adjusting for age and sex determined association between the refined scores in 15 FSHD PBMCs and clinical variables: Ricci, Lamperti scores and disease duration for the refined PAX7 score and LLFF for the refined Lymphoblast score at the 5% significance level.

### Validation of the FSHD muscle-blood biomarker

The 143 gene FSHD muscle-blood biomarker was computed in the 11 control, and 24 TIRM- and matched TIRM+ muscle biopsies and 15 FSHD PBMCs. Comparison between control and FSHD samples was as described for the full PAX7 score. Pearson correlation analysis determined association between the FSHD muscle-blood biomarker in 13 TIRM-FSHD muscle biopsies and matched PBMCs.

The FSHD muscle-blood biomarker was also computed on 26 paired year 1 and year 2 muscle biopsy samples described by Wong et al.,^14^, normalised read counts were obtained from the GEO data base accession: GSE115650. Biomarker levels between year 1 and year 2 samples was compared via a paired Wilcoxon test.

For meta-analysis of the FSHD muscle-blood biomarker as an FSHD biomarker, seven independent datasets containing transcriptomic assessments of FSHD and control muscle biopsies were downloaded as normalized datasets from the GEO database. Rahimov et al.,^55^ GSE36398, describes 50 muscle biopsies assessed by microarray. Bakay et al.,^56^ GSE3307, describes 30 muscle biopsies assessed by microarray. Tasca et al.,^57^ GSE26852, describes 15 muscle biopsies assessed by microarray. Osborne et al.,^43^ GSE10760, describes 49 muscle biopsies assessed by microarray. Dixit et al.,^58^ GSE9397, describes 18 muscle biopsies assessed by microarray. Yao et al.,34 GSE56787, describes 23 muscle biopsies assessed by RNA-seq (control C6 was removed as it was the only non-quadriceps sample). Wang et al.,13 GSE115650, describes 43 muscle biopsies assessed by RNA-seq. These seven datasets describe 228 muscle biopsies (130 FSHD, 98 control). All data were log-transformed and quantile normalised within study for computation of the PAX7 score in line with our previous analyses^9,11,17^. FSHD muscle-blood biomarker differences between FSHD and control samples were evaluated within each study via Wilcoxon test and meta-analysis across the seven studies using a random effects model, with overall significance assessed via Fisher’s combined test.

### Analysis of dataset published by Signorelli et al., 2020

Raw read counts describing RNA-sequencing of PAXgene blood from 54 FSHD patients and 29 controls were described in Signorelli et al., 2020^23^. The dataset comprises 2 independent cohorts processed separately. The Nijmegen cohort comprises 39 FSHD and 11 control samples, and was sequenced to an average depth of 17.5 million reads per sample. The Newcastle cohort comprises 15 FSHD and 18 control samples and was sequenced to an average depth of 43.8 million reads per sample. Clinical annotations included age, sex and Lamperti score. Each dataset was normalised separately using the DESeq2 package in R^59^, data was then log-transformed and quantile normalised within each cohort in line with our prior analysis^10,11,17^. The FSHD muscle-blood biomarker was computed on each sample and the resulting distribution of values *z*-normalised within each cohort and values from the two cohorts combined for subsequent analysis.

Two multivariate linear regression analyses were performed. First, modelling the *z*-normalised FSHD muscle-blood biomarker across all 83 samples as a function of FSHD status, sex, age and cohort. Second, modelling the *z*-normalised FSHD muscle-blood biomarker across the 54 FSHD samples as a function of Lamperti score, sex, age and cohort. Significance of each coefficient was assessed at the 5% level.

The 54 FSHD samples were split into two groups on the basis of age, 26 patients had ages below the cohort mean (49.6 years) and 28 patients above. These subsets were analysed separately, via multivariate regression analyses modelling the FSHD muscle-blood biomarker as a function of mild/severe FSHD (defined as Lamperti score 1-7 for mild, 8-15 for severe), age and sex, significance of each coefficient was assessed at the 5% level. ROC curves and AUC calculations were performed using the ROC package in R^60^ to assess the strength of the FSHD muscle-blood biomarker as a classifier of mild/severe FSHD across all 54 patients and across the 28 older patients separately.

### GSEA of the FSHD muscle-blood biomarker

GSEA was performed separately on the 64 PAX7 up-regulated and 79 PAX7 down-regulated genes comprising the FSHD muscle-blood biomarker, via Fisher’s Exact Test, to determine overlap with gene sets defined by the molecular signatures database^61^.

## Acknowledgments

We acknowledge support of the FSHD Society (FSHS-82016-03 to C.R.S.B. and P.S.Z.) and the Prinses Beatrix Fonds (W.OR15-23 to BvE and LABJ). The Zammit laboratory is funded by the Medical Research Council (MR/P023215/1 and MR/S002472/1), FSHD Society Shack Family and Friends research grant (FSHS-82013-06) and Association Française contre les Myopathies (AFM 17865). Dr. van Engelen reports grants from Global FSH, Stiching Spieren voor Spieren, Dutch FSHD Foundation. We thank Mirko Signorelli and Pietro Spitali for sharing data from Signorelli et al.,^23^ and their input on biomarker analysis.

## Figure and Table legends

**Table S1: *Full clinical annotations of patients and controls employed in the study***. Table displays data for all 49 individuals, whether they underwent PBMC and/or muscle biopsy sampling and which muscles were biopsied, as well as patient age at examination, sex, disease duration and MRC sum score. For FSHD patients MRI assessment of inflammation and fat fractions are displayed as well as Ricci and Lamperti clinical severity scores and MVC of TA, for FSHD1 patients D4Z4 repeat length is displayed.

**Table S2: *Globaltest results assessing the gene sets comprising FSHD biomarkers as correlates of FSHD status***. Table displays results of globaltest implemented to determine association between genes comprising each of the 5 biomarkers (3 DUX4 target gene sets, PAX7 target gene set, Lymphoblast score gene set) and Control/TIRM-/TIRM+ muscle adjusting for age and sex, TIRM-/TIRM+ muscle adjusting for patient, age and sex and Control/FSHD PBMCs adjusting for age and sex.

**Table S3: *Genes comprising the FSHD muscle-blood biomarker***. Table lists genes comprising the FSHD muscle-blood biomarker.

*Table S4: GSEA on the 143 gene FSHD muscle-blood biomarker*. Results of GSEA, via Fisher’s exact test evaluating the 79 PAX7 down regulated targets (Sheet 1) and 64 PAX7 up-regulated targets (Sheet 2) against the gene sets of the Molecular Signatures Database.

